# Identification of *Mycobacterium abscessus* subspecies by MALDI-TOF Mass Spectrometry and Machine Learning

**DOI:** 10.1101/2022.07.28.501950

**Authors:** David Rodríguez-Temporal, Laura Herrera, Fernando Alcaide, Diego Domingo, Neus Vila, Manuel J. Arroyo, Gema Méndez, Patricia Muñoz, Luis Mancera, María Jesús Ruiz-Serrano, Belén Rodríguez-Sánchez

## Abstract

*Mycobacterium abscessus* complex is one of the most common and pathogenic nontuberculous mycobacteria (NTM) isolated in clinical laboratories. It consists of three subspecies: *M. abscessus* subsp. *abscessus, M. abscessus* subsp. *bolletii* and *M. abscessus* subsp. *massiliense*. Due to their different antibiotic susceptibility pattern, a rapid and accurate identification method is necessary for their differentiation. Although matrix assisted laser desorption/ionization-time of flight mass spectrometry (MALDI-TOF MS) has proven useful for NTM identification, the differentiation of *M. abscessus* subspecies is challenging. In this study, a collection of 244 clinical isolates of *M. abscessus* complex was used for MALDI-TOF MS analysis and for the development of machine learning predictive models. Overall, using a Random Forest model with several confidence criteria (samples by triplicate and similarity values >60%), a total of 95.8% of isolates were correctly identified at subspecies level. In addition, differences in culture media, colony morphology and geographic origin of the strains were evaluated, showing that the latter most affected the mass spectra of isolates. Finally, after studying all protein peaks previously reported for this complex, two novel peaks with potential for subspecies differentiation were found. Therefore, machine learning methodology has proven to be a promising approach for rapid and accurate identification of subspecies of the *M. abscessus* complex using MALDI-TOF MS.

## INTRODUCTION

Nontuberculous mycobacteria (NTM) are a group of mycobacteria present in the environment that, in some cases, can cause different types of infections in humans, such as pulmonary infections, skin and soft tissue infections and disseminated infections (1). *Mycobacterium abscessus* complex is one of the most common and pathogenic NTM isolated in clinical laboratories, causing respiratory infections, even in patients with cystic fibrosis (2). *M. abscessus* complex contains three subspecies: *M. abscessus* subsp. *abscessus, M. abscessus* subsp. *massiliense* and *M. abscessus* subsp. *bolletii* (3). Hereafter, they will be referred as *M. abscessus, M. massiliense* and *M. bolletii*, respectively.

The three subspecies show different susceptibility to clarithromycin, a decisive antibiotic for the treatment of these infections. Thus, *M. bolletii* and most strains of *M. abscessus* shows resistance to clarithromycin, whereas *M. massiliense* is susceptible (4). The different antibiotic susceptibility pattern, in addition to the recommendation of the American Thoracic Society and Infectious Diseases Society of America (ATS/IDSA)to identify NTMs at species level, makes it necessary to implement novel approaches for rapid and accurate discrimination of these three subspecies (5).

Currently, *M. abscessus* complex subspecies can only be identified by molecular methods, such as a commercial kit based on PCR-reverse hybridization (6) or by multiple gene sequencing (*hsp65, rpoB, erm*(41), etc.) (7, 8). On the other hand, the use of Matrix Assisted Laser Desorption/Ionization-Time of Flight Mass Spectrometry (MALDI-TOF MS) allows the reliable identification of most NTMs and has become the main identification method in several clinical laboratories (9, 10). However, differentiation of closely related species (like *M. abscessus* complex subspecies) remains a challenge. Although some studies have attempted subspecies identification by protein peak analysis (11–16), there is no consensus on the best strategy to follow. In last years, new approaches for data analysis from MALDI-TOF mass spectra have been applied, such as machine learning methods, which has the potential to get additional information than simple species identification (17).

The aim of this study was to evaluate MALDI-TOF MS and Machine Learning algorithms for the differentiation of *M. abscessus* complex subspecies. This study represents the first proof of concept for the identification of these species by applying MALDI-TOF MS and Machine Learning.

## MATERIALS & METHODS

### Mycobacterial isolates

A total of 244 clinical isolates of *M. abscessus* complex obtained from 152 different patients were included in this study. They encompassed 119 *M. abscessus*, 84 *M. massiliense* and 41 *M. bolletii* isolates. All the isolates were obtained from Hospital General Universitario Gregorio Marañón (HGM; Madrid, Spain), Hospital Universitario La Princesa (HLP; Madrid, Spain), Instituto de Salud Carlos III-Centro Nacional de Microbiología (ISCIII; Majadahonda, Spain) and Hospital Universitari de Bellvitge (HUB; Hospitalet de Llobregat, Spain). All isolates are described in Table S1.

### Bacterial cultures and protein extraction procedure

All isolates were previously identified by PCR-reverse hybridization (GenoType NTM-DR, Hain Lifescience, Nehren, Germany) or whole genome sequencing. All HGM, HLP and ISCIII isolates were cultured from frozen stocks on 7H11 agar plates until growth was observed. Among HUB isolates, 38 were cultured on 7H11 agar plates and 48 on Löwenstein-Jensen (BioMérieux, Marcy l’Etoile, France) media. In all cases, the isolates were incubated at 37°C until growth was observed (4-7 days). The protein extraction procedure for MALDI-TOF MS analysis was performed as previously described (10). First, a 1 μl loopful of biomass was suspended in 300 μl of High-Pressure Liquid Chromatography (HPLC) quality water, and then heat inactivated in a dry bath at 95°C during 30 min. After this, 900 μl of ethanol were added, the tubes were centrifuged at 13,000 rpm for 2 min and the supernatant was discarded. After centrifuge and discard the supernatant again, the pellet was dried at room temperature. Then, 0.5 mm silica/zirconia beads were added together with 10 μl of acetonitrile. The tubes were vortexed briefly and sonicated for 15 min. After sonication, 10 μl of formic acid were added, the tubes were vortexed for 10 s and centrifuged at 13,000 rpm for 2 min. One microliter of the supernatant was deposited onto the MALDI target plate (Bruker Daltonics, Bremen, Germany) in triplicates, allowed to dry and covered with 1 μl of α-cyano-4-hydroxycinnamic acid (HCCA).

### Spectra acquisition by MALDI-TOF MS and data processing

Acquisition of protein spectra was performed using the MBT Smart MALDI Biotyper (Bruker Daltonics) in the range of 2,000-20,000 Da. All spots were read three times, resulting in 9 protein spectra per isolate. The spectra were exported and processed with Clover MS Data Analysis software (Clover Biosoft, Granada, Spain). The processing pipeline consisted on: 1) Smoothing by Savitzky-Golay filter (window length=11, polynomial order=3); 2) Baseline substraction by Top-Hat filter (factor=0.02); 3) Alignment of spectra with 2 Da of constant tolerance and 300 ppm of linear mass tolerance; and 4) Normalization by Total Ion Current (TIC).

### Predictive models and external validation

Once the spectra were processed, unsupervised −Principal Component Analysis (PCA), Hierarchical Cluster Analysis (HCA)- and supervised −Partial Least Squares Discriminant Analysis (PLS-DA), Support Vector Machine (SVM), Random Forest (RF) and K-Nearest Neighbors (KNN)- algorithms were applied for the creation of predictive models. A total of 43 isolates (20 *M. abscessus*, 15 *M. massiliense* and 8 *M. bolletii*) collected in HGM, ISCIII and HUB were included in the test set for the creation of the predictive models, they represented a total of 539 mass spectra. These isolates were randomly selected in order to represent all the variability observed previously (subspecies, morphology, culture media and geographical origin). *M. massiliense* and *M. bolletii* spectra were balanced by oversampling in order to obtain the same number of spectra for each category. Internal validation was performed by 10-fold cross validation. For external validation, 201 isolates collected in all centres were used (99 *M. abscessus*, 69 *M. massiliense* and 33 *M. bolletii*), and the identification obtained in each of the three spots used was considered.

### Ethics statement

The Ethics Committee of the Gregorio Marañón Hospital (CEIm) evaluated this project and considered that all the conditions for waiving informed consent were met, since the study was conducted with microbiological samples and not with human products.

## RESULTS

### Analysis of isolates by unsupervised algorithms

After the analysis of all isolates by PCA, two main clusters were observed (Figure 1). Different variables included in the study that may influence on the mass spectra were examined: the *M. abscessus* complex subspecies, the morphology of the colonies, the type of culture media and the geographical origin of the isolates. As can be observed in Figure 1, the two main clusters corresponded to the geographical origin of the isolates, separating those collected in Madrid hospitals (HGM, HLP and ISCII) from those collected in Barcelona (HUB). On the other hand, isolates from different subspecies and different morphology overlapped in both clusters, as well as isolates from HUB, which were analyzed in two different culture media.

**Figure 1.**
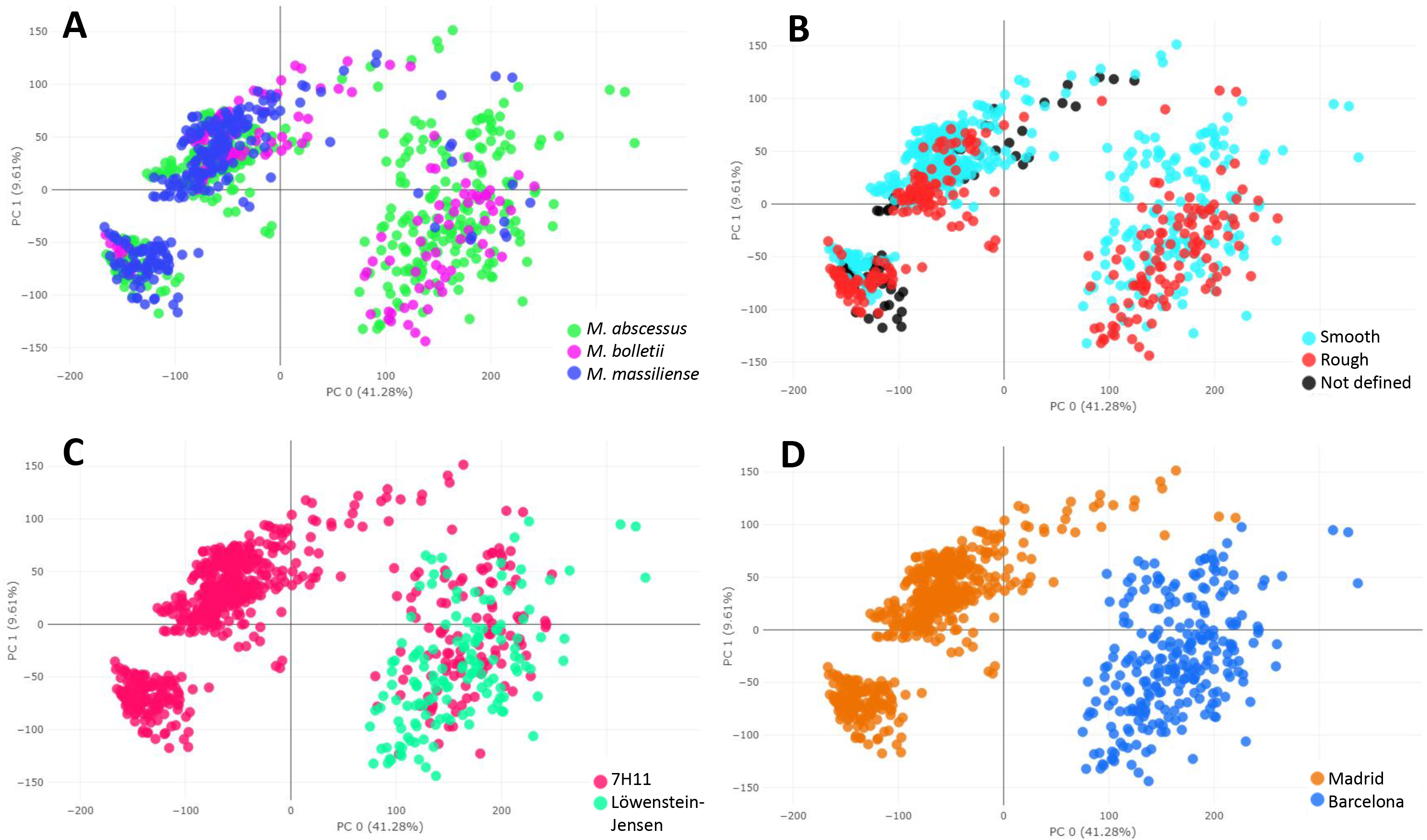
Principal Component Analysis of all isolates included in the study, colored according to different characteristics: **A**, comparison of *M. abscessus* complex subspecies; **B**, comparison of colony morphology; **C**, comparison of culture media; **D**, comparison of geographical zone of origin.

### Analysis of isolates by supervised algorithms

#### Internal validation of predictive models

The results for each algorithm after applying a 10-fold cross validation are showed in Table 1. The algorithms PLS-DA (Figure S1), SVM and RF showed the same accuracy (99.8%), with only one spectrum of the 539 misclassified as other category (Table S2), while KNN was the algorithm with lower accuracy (Figure S1).

**Table 1.**
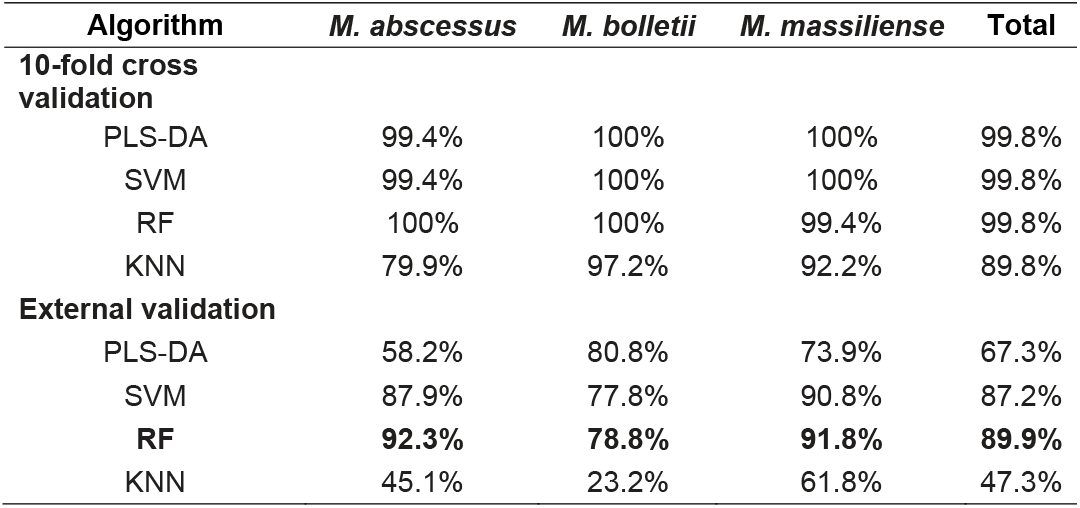
Accuracy results for internal 10-fold cross validation and external validation.

| Algorithm | *M. abscessus* | *M. bolletii* | *M. massiliense* | Total |
| --- | --- | --- | --- | --- |
| **10-fold cross validation** |  |  |  |  |
| PLS-DA | 99.4% | 100% | 100% | 99.8% |
| SVM | 99.4% | 100% | 100% | 99.8% |
| RF | 100% | 100% | 99.4% | 99.8% |
| KNN | 79.9% | 97.2% | 92.2% | 89.8% |
| **External validation** |  |  |  |  |
| PLS-DA | 58.2% | 80.8% | 73.9% | 67.3% |
| SVM | 87.9% | 77.8% | 90.8% | 87.2% |
| **RF** | **92.3%** | **78.8%** | **91.8%** | **89.9%** |
| KNN | 45.1% | 23.2% | 61.8% | 47.3% |

#### External validation of predictive models

Blind analysis of the 201 isolates used for external validation showed that PLS-DA and KNN produced low identification rates, while RF algorithm yielded 89.9% correct classification (Table 1). With this algorithm, *M. bolletii* obtained the lower identification rate, with 21 spectra misclassified (Table S3). Due to RF obtained the highest identification rate (Figure 2A), its results were further analyzed. Among the three identifications obtained in each spot for each isolate, the subspecies obtained in at least 2 spots was considered as the final identification. A total of 184 (91.5%) of the isolates obtained uniform identification results (Table 2), and only one isolate obtained different subspecies identification for each spot. Among the isolates with identical identification, the accuracy rate of subspecies level identification was higher than in those with only two matching identifications. Moreover, the probability of correct identification provided by RF was evaluated in order to establish a confidence cut-off. For all subspecies, 172 (85.6%) isolates obtained a probability higher than 60% (Figure 2B), so this cut-off was proposed for confident result. Considering the categorical result, 89.6% of isolates were correctly identified at subspecies level, while establishing the confidence cut-off at 60% of probability, the accuracy rate increased to 95.3% (Figure 2C). Moreover, when both parameters were considered (same identification in three spots and confidence higher than 60%), out of the 170 isolates that met these criteria, 163 (95.8%) were correctly identified (Table 2). The three subspecies performed similar, with similar Area Under the Curve (AUC) between them (Figure 2D). Finally, Positive Predictive Values (PPV) were evaluated for each subspecies. Considering all identification results, the PPV obtained was 89.3% for *M. abscessus*, 89.3% for *M. bolletii* and 90.0% for *M. massiliense*. When we considered only those isolates with a probability result higher than 60%, the PPV increased to 96.4%, 91.7% and 95.3% for each subspecies, respectively (Figure 2E).

**Figure 2.**
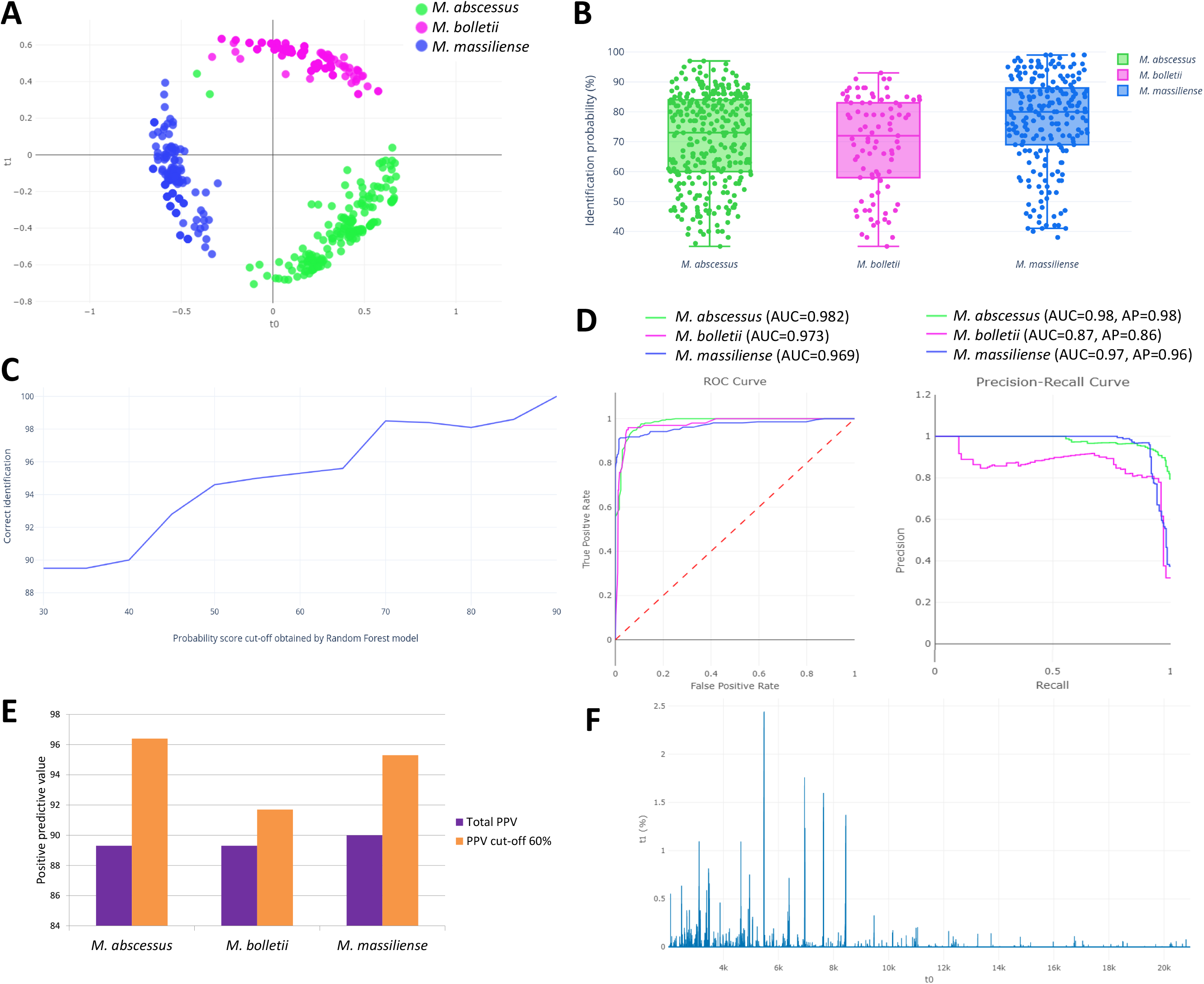
Analysis of mass spectra by Random Forest (RF) algorithm. **A.** RF plot of the model. **B.** Percentages of identification probably obtained by RF on validation isolates. **C.** Number of correctly identified isolates according to probability cut-off obtained by RF. **D.** ROC and Precision Recall curves for validation isolates by RF. **E.** Total Positive Predictive Value (PPV) for RF results and PPV using a 60% probability cut-off. **F.** Feature importances of mass peaks for RF model.

**Table 2.** Random Forest (RF) accuracy according to the identification (ID) obtained in each spot.

| RF identification | N isolates (%) | N correct (%) |
| --- | --- | --- |
| Same ID in 3 spots | 184 (91.5%) | 173 (94.0%) |
| Same ID in 2 spots | 16 (8.0%) | 7 (43.8%) |
| Three different ID | 1 (0.5%) | 0 (0%) |
| Isolates with confidence >60% | 172 (85.6%) | 164 (95.3%) |
| **Same ID (3 spots) and confidence >60%** | **170 (84.6%)** | **163 (95.8%)** |

### Specific peak analysis for subspecies discrimination

All protein peaks reported in previous studies were searched among the analyzed isolates (Table 3). No unique subspecies specific peaks were found, although some of them were present in most strains of certain subspecies. Thus, almost all *M. abscessus* isolates showed peaks at 2081, 3378 and 7637 *m/z; M. bolletii* showed 2081, 3123, 3463 and 7637 *m/z;* and *M. massiliense* showed 3378, 4385 and 6711 *m/z*. In addition, two novel potential peaks were found in this study: 2673 *m/z*, which was present in 88.9% of *M. abscessus* isolates, 17.1% of *M. bolletii* and 7.3% of *M. massiliense;* and 6960 *m/z*, which was present in 90.5% of *M. abscessus* isolates, 9.8% of *M. bolletii* and 26.0% of *M. massiliense* (Table 3).

**Table 3.** Presence of all protein peaks reported in previous studies and in the present one among isolates of each subspecies. NA: not analyzed.

| *m/z* | *M. abscessus* |  | *M. bolletii* |  | *M. massiliense* |  |
| --- | --- | --- | --- | --- | --- | --- |
|  | Present study (%) | Previous studies (%) | Present study (%) | Previous studies (%) | Present study (%) | Previous studies (%) |
| **Previously reported peaks** |  |  |  |  |  |  |
| 2081 | 96.0 | 31.7-96.5 | 100 | 100 | 53.1 | 0 |
| 3108 | 10.3 | 0 | 17.1 | 0 | 79.2 | 100 |
| 3123 | 84.1 | 100 | 95.1 | 100 | 9.4 | 0 |
| 3354 | 4.0 | 1.4 | 12.2 | NA | 3.1 | 100 |
| 3378 | 99.2 | 100 | 36.6 | 0 | 97.9 | 100 |
| 3463 | 23.8 | 0 | 90.2 | 66.7 | 31.2 | 0 |
| 4385 | 31.7 | 0-1.4 | 29.3 | 0 | 89.6 | 89.5-100 |
| 4391 | 53.2 | 98.6-100 | 41.5 | 100 | 9.4 | 0-5.2 |
| 6711 | 32.5 | 0 | 73.2 | NA | 90.6 | 100 |
| 7637 | 92.6 | 93.2-100 | 90.2 | 100 | 62.5 | 0 |
| 7667 | 11.9 | 3.4 | 2.4 | 0 | 41.7 | 88.1-100 |
| 8508 | 7.1 | 4.9 | 14.6 | NA | 9.4 | 84.6 |
| 8768 | 2.4 | 0-0.7 | 7.3 | 0 | 70.8 | 38.4-100 |
| 8782 | 72.2 | 89.2-100 | 61.0 | 100 | 6.2 | 0-2.8 |
| 9475 | 78.6 | 17-100 | 73.2 | 100 | 16.7 | 0-9.5 |
| **Novel potential peaks** |  |  |  |  |  |  |
| 2673 | 88.9 | NA | 17.1 | NA | 7.3 | NA |
| 6960 | 90.5 | NA | 9.8 | NA | 26.0 | NA |

## DISCUSSION

Differentiation of *M. abscessus* complex subspecies by MALDI-TOF MS has been attempted in previous studies using conventional peak analysis (11–16). However, many variables can hinder this objective, such as the culture media used, the morphology of the colonies or the geographic origin of the strains (14, 18). Moreover, due to the large number of protein peaks that are usually found in mass spectra, accurate identification based on only a few peaks may not be entirely reliable. Therefore, it is necessary to apply novel strategies capable of analyzing a large amount of data, such as machine learning methodologies (17, 19).

In the present study, we applied machine learning using both unsupervised and supervised algorithms. The first approach by unsupervised methodology (PCA) did not provide subspecies differentiation (Figure 1). In the case of morphology, no important spectral differences between smooth and rough variants were observed. The main difference of these two variants is the expression of glycopeptidolipids on the surface (20, 21), and due to MALDI-TOF MS analyzes mainly ribosomal proteins, these differences were not detected. On the other hand, because this is a proof of concept and identification of mycobacteria from liquid media could be more complex (22), only solid culture media were evaluated, and differences between 7H11 and Löwenstein-Jensen were not observed. Interestingly, the two main clusters obtained by PCA corresponded to the geographic origin of the isolates. All strains obtained from the three Madrid hospitals (HGM, HLP and ISCIII) grouped together, while those from Barcelona (HUB) were separated. All isolates from Madrid were analyzed in the same hospital (HGM) and Barcelona isolates in HUB by the same operator, so differences in experience and preparation of the protein extracts were discarded. In addition, the MALDI-TOF MS model in both centers was the same (MBT Smart Biotyper) and the acquisition of spectra was performed with the same technical parameters, so the influence of the instruments was minimal. Therefore, these results may suggest that differences in mass spectra of *M. abscessus* complex from different origins is greater than expected, and highlights the importance of include strains from diverse origins in this type of studies.

The application of supervised machine learning algorithms was targeted to differentiation of the three subspecies. Among the four algorithms tested, the lower results were obtained by PLS-DA (Figure S2A), SVM (Figure S2B) and KNN (Figure S2C), while Random Forest was able to identify a greater number of isolates at subspecies level (Table 1). As recommended by other studies, the identification of NTM by MALDI-TOF MS should be performed in 2 or 3 replicates (10), so for more accurate identifications we used three spots for each isolate. When the identification of the three spots was considered, the accuracy was higher in those cases where the same subspecies was obtained in all spots (Table 2). The categorical result of RF is accompanied by a probability result, so we aimed to establish a confidence cut-off in order to reach higher accuracy of identification. Without applying probability cut-off, RF correctly identified 89.9% of the isolates. Most of isolates (172; 85.6%) obtained probability results above 60% (Figure 2B), so when the cut-off was established at 60%, the accuracy rate increased to 95.3% (Figure 2C). Moreover, by applying this cut-off, the PPV for all subspecies increased to higher than 90% (Figure 2E). Among the 170 isolates that met the criteria of obtaining the same identification in all spots and a confidence higher than 60%, a total of 95.8% were correctly classified, with 7 isolates misidentified: 2 *M. abscessus* identified as *M. massiliense*, 2 *M. massiliense* identified as *M. abscessus*, 2 *M. massiliense* as *M. bolletii* and 1 *M. bolletii* identified as *M. abscessus*.

In order to analyze the protein peaks found in this study, all previously reported peaks were searched and compared with our isolates. In most cases, the detection rate of the peaks was similar to previous reports (Table 3) and, in addition, the most important peaks were found in the range of 2000-10000 Da (Figure 2F). However, remarkable discrepancies in few cases were found. Some peaks were found with a lower presence than reported previously: this is the case of the 4391 m/z peak (Figure S3A) in *M. abscessus* and *M. bolletii*, with only half of our isolates presenting it; the peak around 8782 m/z (Figure S3B) that was present in 61% of *M. bolletii* isolates in comparison with 100% reported by Suzuki et al. (12) and Kehrmann et al. (15); and the peak around 7667 m/z (Figure S3C) in *M. massiliense* that was found in 47.1% of our isolates. Strikingly, peaks 3354 and 8508 m/z were found only in a few *M. massiliense* isolates, while they were previously reported in most isolates of this subspecies (13, 14). On the other hand, peaks that were reported as absent in some subspecies, were found in some of our isolates. That was the case of 3108 (Figure S3D) and 4385 m/z (Figure S3A) in *M. abscessus* and *M. bolletii*; 3123 m/z (Figure S3D) in *M. massiliense*; 3378 m/z in *M. bolletii* (Figure 3A); 3463 m/z (Figure S3E) in *M. abscessus* and *M. massiliense*; and 6711 m/z (Figure S3F) in *M. abscessus*. The greater differences were in peaks 2081 (Figure 3B) and 7637 m/z, which have never been reported in *M. massiliense* (12, 16) and we found them in more than 50% of *M. massiliense* isolates. All these differences could have been influenced by two factors. First, it is important to include a high number of strains, representing the three subspecies in order to confirm that the peaks found are specific to them. The second factor is the geographic origin of the isolates. There have been reported differences in peak patterns according to the origin of the strains (14), so multicentric studies are needed to search common peaks worldwide and create accurate identification algorithms. On the other hand, two novel potential peaks have been found: 2673 (Figure 3C) and 6960 m/z (Figure 3D), both of them present in most *M. abscessus* isolates and in low number of isolates from the others subspecies.

**Figure 3.**
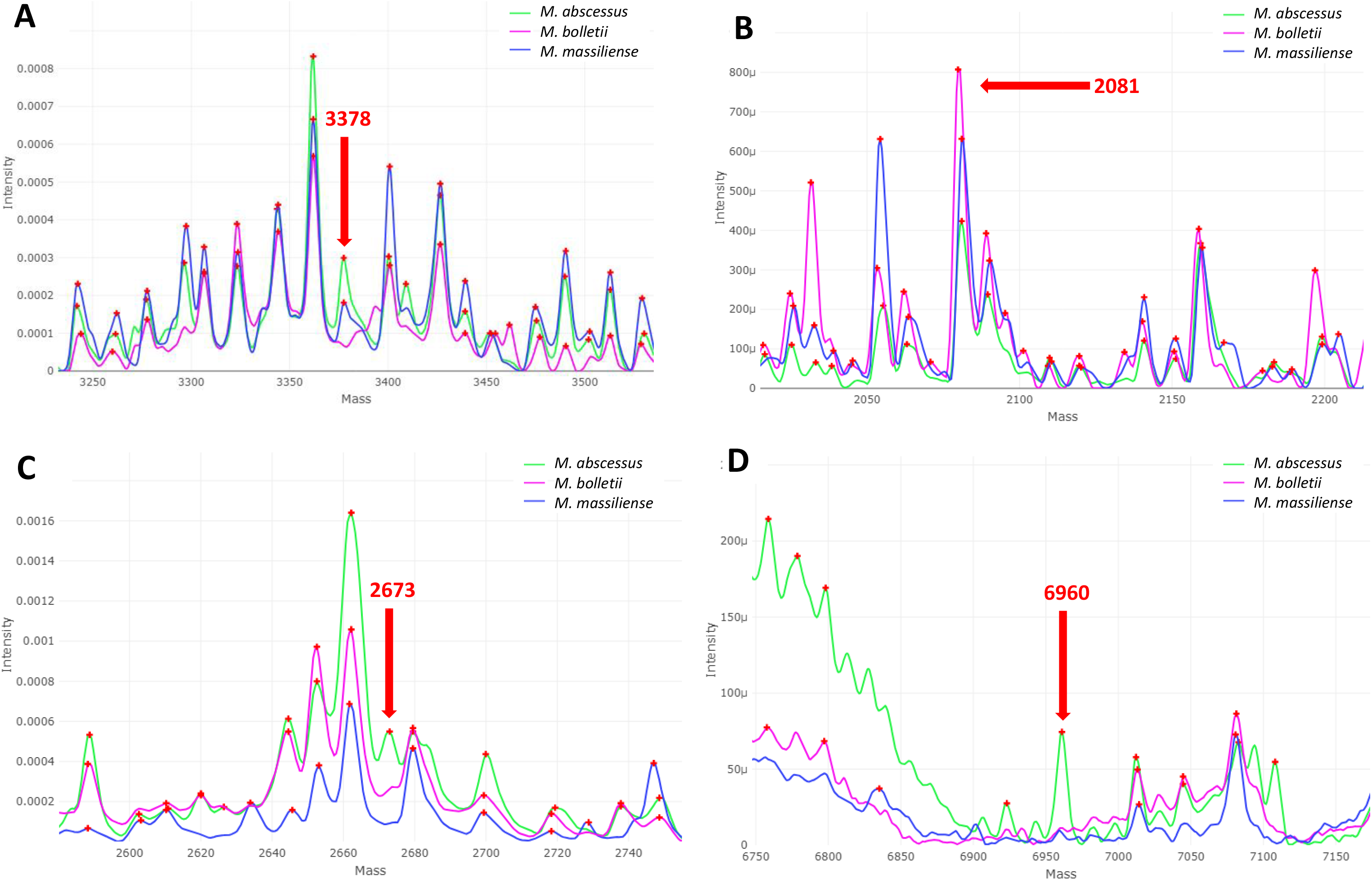
Novel potential protein peaks reported in the present study and other relevant peaks. **A.** 3,378 m/z. **B.** 2,081 m/z. **C.** 2,673 m/z. **D.** 6,960 m/z.

Due to the variability in the detection of peaks observed previously, the present study showed that the application of novel methodologies for data analysis, such as machine learning, could be an innovative way for improve the MALDI-TOF MS accuracy on identifying *M. abscessus* subspecies. Recently, other novel strategies have been evaluated for the same purpose. Khor et al. used the MALDI Biotyper Sirius system (Bruker Daltonics) for the detection of subspecies-specific lipids, and was able to differentiate a few *M. abscessus* complex isolates (23). On the other hand, Bajaj et al. evaluated for the first time the Liquid Chromatography-Mass Spectrometry for identification of *M. abscessus* subspecies (24). However, these novel methods need to be validated with larger collections of clinical isolates to confirm their utility in a microbiology laboratory setting.

In conclusion, the high correct identification rate of *M. abscessus* complex subspecies obtained in this study, states the utility of machine learning strategy for identification purposes. This method could be further refined in near future by the addition of a greater number and diversity of isolates.

## Author contributions

DRT: conceptualization, experimentation, formal analysis, data collection, validation, visualization, original draft preparation and review/editing. LH, FA, DD, NV, PM, MJRS: submission of isolates, writing and review/editing. MJA GM, LM: data analysis, validation, writing and review/editing. BRS: conceptualization, project administration, formal analysis, supervision, validation, visualization, original draft preparation and review/editing.

